# Large-scale Lassa fever outbreaks in Nigeria: quantifying the association between disease reproduction number and local rainfall

**DOI:** 10.1101/602706

**Authors:** Shi Zhao, Salihu S. Musa, Hao Fu, Daihai He, Jing Qin

**Author notes:** Correspondence to (S.Z.) & (D.H.).

## Abstract

**Background:** Lassa fever (LF) is increasingly recognized as an important rodent-borne viral hemorrhagic fever presenting a severe public health threat to sub-Saharan West Africa. In 2018, LF caused an unprecedented outbreak in Nigeria, and the situation was worse in 2019. This work aims to study the epidemiological features of outbreaks in different Nigerian regions and quantify the association between reproduction number (*R*) and local rainfall by using modeling analysis.

**Methods:** We quantify the infectivity of LF by the reproduction numbers estimated from four different growth models: the Richards, three-parameter logistic, Gompertz, and Weibull growth models. LF surveillance data are used to fit the growth models and estimate the *R*s and epidemic turning points (*τ*) in different regions at different time periods. Cochran’s Q test is further applied to test the spatial heterogeneity of the LF epidemics. A linear random-effect regression model is adopted to quantify the association between *R* and local rainfall with various lag terms.

**Findings:** Our estimated *R*s for 2017-18 (1.33 with 95% CI: [1.29, 1.37]) and 2018-19 (1.29 with 95% CI: [1.27, 1.32]) are significantly higher than those for 2016-17 (1.23 with 95% CI: [1.22, 1.24]). We report spatial heterogeneity in the *R*s for outbreaks in different Nigerian regions. For the association between rainfall and *R*, we find that a one unit (mm) increase in average rainfall over the past 7 months could cause a 0.62% (95% CI: [0.20%, 1.05%]) rise in *R*.

**Conclusion:** There is significant spatial heterogeneity in the LF epidemics in different Nigerian regions. We report clear evidence of rainfall impacts on LF outbreaks in Nigeria and quantify the impact.

## Introduction

Lassa fever (LF), caused by Lassa virus (LASV), is increasingly recognized as an important rodent-borne viral hemorrhagic fever presenting a severe public health threat to some of the communities in sub-Saharan West Africa [1]. Discovered in 1969 [2], LF is endemic to much of rural Nigeria and regions in the Mano River Union [3]. LASV transmits from human to human, as well as via the zoonotic cycle [1, 3, 4]. LF has a high case fatality rate ranging from 1% in the community to over 60% in hospital settings [1, 4, 5]. The common reservoir of LASV is *Mastomys natalensis*, one of the most widespread rodent species in sub-Saharan Africa [1, 3], which exhibit sensitive population dynamics to water level [6, 7]. Previous studies have recognized the ecological connection between the population levels of rodents and rainfall [8–10].

LF epidemics typically start in November and last until May of the following year, with the majority of cases occurring in the first quarter of the following year, in addition to sporadic cases reported throughout the year. The 2017-18 epidemic in Nigeria was an unprecedented LF outbreak in the country’s history [11], which resulted in 400 confirmed cases, including 97 deaths, between January and March 2018 [12]. The 2018-19 epidemic in Nigeria has already caused 420 confirmed cases from January to March 02 of 2019, which included 93 deaths [12]. The five states of Edo, Ondo, Ebonyi, Bauchi, and Plateau are the only states that have been among the top 10 hit hardest states in terms of number of LF cases in both the 2018 (85.5% of total national cases) and 2019 (85.7% of total national cases) outbreaks. While there have been discussions about the connection of rainfall to LF [13, 14], this connection has not yet been demonstrated and quantified. This work aims to study the epidemiological features of outbreaks in different Nigerian regions between January 2016 and March 2019. We estimate LF infectivity in terms of the reproduction number (*R*) and quantify the association between *R* and local rainfall by using modeling analysis. We explore the spatial heterogeneity of the LF outbreaks and summarize the overall findings with model-average estimates.

## Data and methods

### Data

Weekly Lassa fever (LF) surveillance data are obtained from the Nigeria Centre for Disease Control (NCDC) [12]. Laboratory-confirmed case time series are used for analysis. We examine the major epidemics that occurred between January 2016 and March 2019 across the whole country and only in the five states that were among the top 10 hardest-hit states in both the 2018 and 2019 outbreaks, i.e., Edo, Ondo, Ebonyi, Bauchi, and Plateau. Local rainfall records are collected from the historical records of the World Weather Online website [15]. Figure 1(a)-(b) shows the rainfall time series of the five states and the weekly reported LF cases across the entirety of Nigeria.

**Figure 1.**
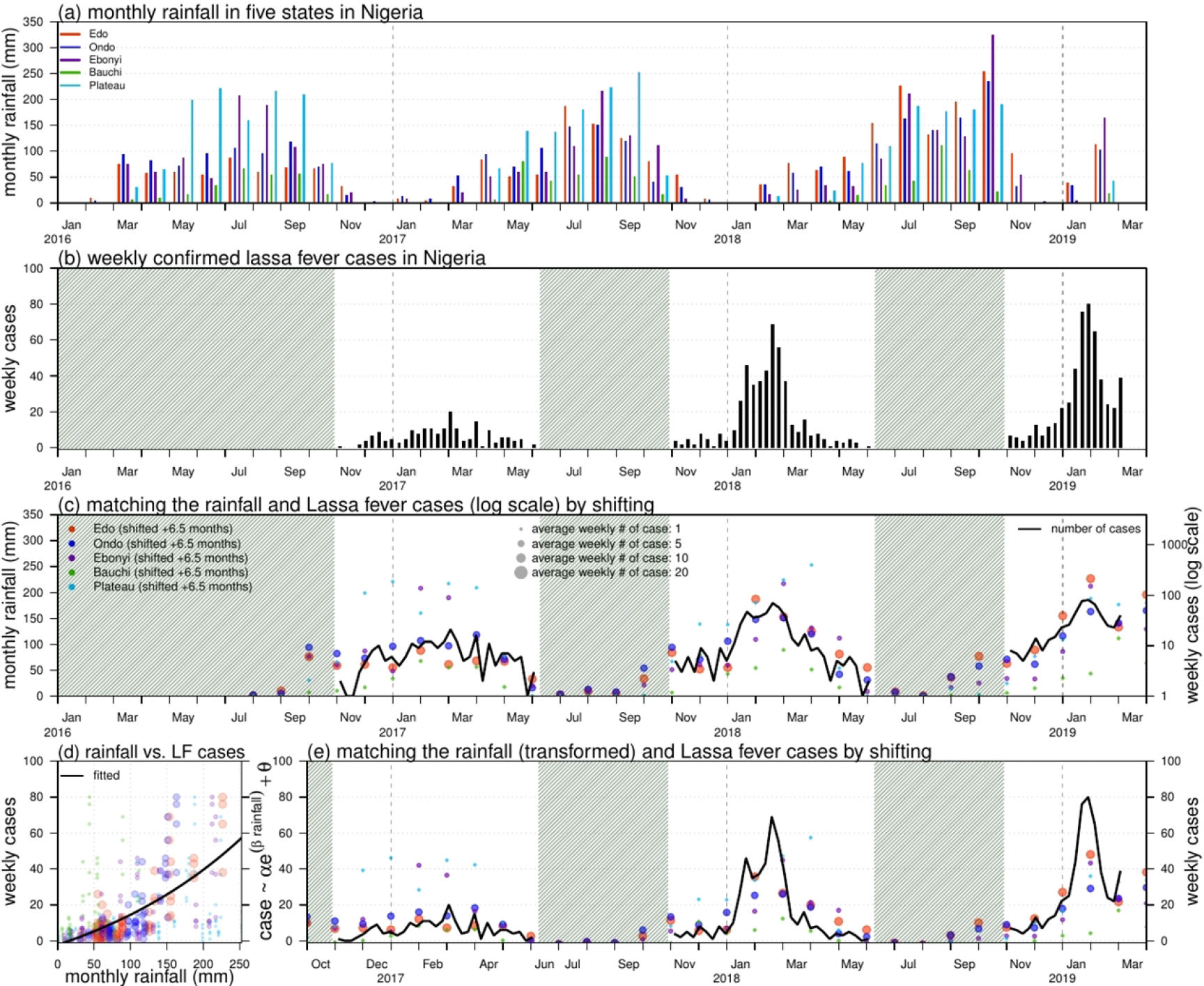
Rainfall (unit: mm) and number of Lassa fever (LF) cases in Nigeria. Panel (a) shows the monthly rainfall in five states in Nigeria. Panel (b) shows the weekly number of LF cases in Nigeria. The shaded area represents weekly number of cases lower than 10. Panel (c) matches the rainfall (dots) and LF cases (in log scale, black line) by shifting the rainfall time series by +6.5 months. The sizes of each dot represent the number of the average weekly LF cases in each state in the 2017-18 and 2018-19 outbreaks. Panel (d) is the scatter plot of rainfall (shifted +6.5 months) versus LF cases; the dots of different colors and sizes share the same scheme as in panel (c). The black line is the fitting outcome of the formula “case ~ *α* exp(*β* * rainfall) + *θ*” by least square estimation; the fitting R-squared is 0.43 and significance is *p*-value < 0.0001. Panel (e) is the fitting outcome from panel (d), and the rainfall dots (shifted +6.5 months) of different colors and sizes share the same scheme as in panel (c).

### Models and estimation

Four different nonlinear growth models are adopted to pinpoint the epidemiological features of each outbreak. The models are the Richards, three-parameter logistic, Gompertz, and Weibull growth models. These simple structured models are widely used to study S-shaped cumulative growth processes; e.g., the curve of a single-wave epidemic and have been extensively studied in previous work [16, 17]. These models consider cumulative cases with saturation in the growth rate to reflect the progression of an epidemic due to reductions in susceptible pools. The extrinsic growth rate does not steadily decline but increases to a maximum (i.e., saturation) before steadily declining to zero.

We fit all models to the weekly reported LF cases in different regions evaluate the fitting performance by the Akaike information criterion (AIC). We adopt the standard nonlinear least squares (NLS) approach for model fitting and parameter estimation, following [16, 18]. A *p*-value < 0.05 is regarded as statistically significant, and the 95% confidence intervals (CIs) are estimated for all unknown parameters. The models are selected by comparing the AIC to that of the baseline model. Only the models with an AIC lower than the AIC of the baseline model are considered for further analysis. Since the epidemic curves of an infectious disease commonly exhibit autocorrelations [19], we use autoregression (AR) models with a degree of 2, i.e., AR(2), as the baseline models for growth model selection. We also adopt the coefficient of determination (R-squared) and the coefficient of partial determination (partial R-squared) to evaluate goodness-of-fit and fitting improvement, respectively. For partial R-squared, the AR(2) model is treated as the baseline model.

After the selection of models, we estimate the epidemiological features (parameters) of turning point (*τ*) and reproduction number (*R*) from the selected models. The turning point is defined as the time point of a sign change in the rate of case accumulation, i.e., from increasing to decreasing or *vice versa* [16, 18]. The reproduction number, *R*, is the average number of secondary infectious cases produced by one infectious case during a disease outbreak [18, 20]. When the population is totally (i.e., 100%) susceptible, the *R* will equate to the basic reproduction number, commonly denoted as *R_0_* [20, 21]. The reproduction number (*R*) is given in Eqn (1),

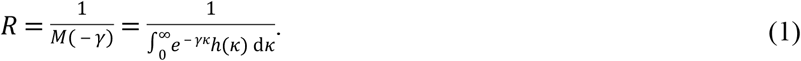

Here, *γ* is the intrinsic per capita growth rate from the nonlinear growth models, and *κ* is the serial interval of the Lassa virus (LASV) infection. The serial interval (i.e., the generation interval) is the time between successive cases in a chain of transmission [20, 22, 23]. The function *h*(⋅) represents the probability distribution of *κ*. Hence, the function *M*(⋅) is the Laplace transform of *h*(⋅), and specifically, *M*(⋅) is the moment generating function (MGF) of a probability distribution [20]. According to previous work [24], we assume *h*(*κ*) to follow a Gamma distribution with a mean of 7.8 days and a standard deviation (SD) of 10.7 days. Therefore, *R* can be estimated with the values of *γ* from the fitted models [18, 20, 25].

We then summarize the *κ* and *R* estimates by AIC-weighted model averaging. The AIC weights, *w*, of the selected models (with AIC lower than the AIC from the AR(2) model) are defined in Eqn (2),

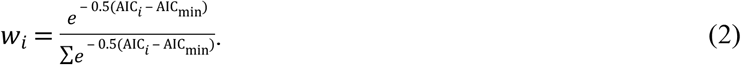

Here, AIC_*i*_ is the AIC of the *i*-th selected model, and the AIC_min_ is the lowest AIC among all selected models. Thus, the *i*-th selected model has a weight of *w*_*i*_. The model-averaged estimator is the weighted average of the estimates in each selected model, which has been well studied in previous work [16, 26].

### Testing the spatial heterogeneity of the LF epidemics

After finding the model-averaged estimates, we apply Cochran’s Q test to examine the spatial heterogeneity of the epidemics in different regions over the same period of time [27]. For instance, we treat the model-averaged *R* estimates as the univariate meta-analytical response against different Nigerian regions (states) and further check the heterogeneity by estimating the significance levels of the Q statistics. A *p*-value < 0.05 is regarded as statistically significant.

### Association between rainfall and reproduction number

The association between local rainfall and LASV transmissibility are modeled by a linear random-effect regression (LRER) model in Eqn (3),

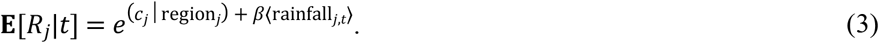

Here, **E**(⋅) represents the expectation function and *j* is the region index corresponding to different regions (states). Term *c*_*j*_ is the interception term of the *j*-th region to be estimated and it is variable from different regions, serving as the baseline scale of transmissibility in different states. The term *t* denotes the cumulative lag in the model, and 〈rainfall_*j,t*_〉 represents the average monthly rainfall of the previous *t* months from the turning point, *τ*, of the *j*-th region. The reproduction numbers, *R*_*j*_, are estimated for different epidemics from the selected growth models. The regression coefficient, *β*, is to be estimated. Hence, the term (*e^β^* - 1)*100% is the percentage change rate (of *R*), which can be interpreted as the percentage change in transmissibility due to a one unit (mm) increase in monthly rainfall. The framework of the regression is based on the exponential form of the predictor to model the expectation of transmissibility (e.g., *R*); this framework is inspired by previous work [28–31]. To quantify the impacts of local rainfall, we calculate the percentage change rate with different cumulative lags (*t*) from 0-11 months and estimate their significant levels. Only the lag terms (*t*) with significant estimates are presented in this work.

We present the modeling analysis procedures in a flow diagram in Figure 2. All analyses are conducted by using R (version 3.4.3 [32]), and the R function “nls” is employed for the NLS estimation of model parameters.

**Figure 2.**
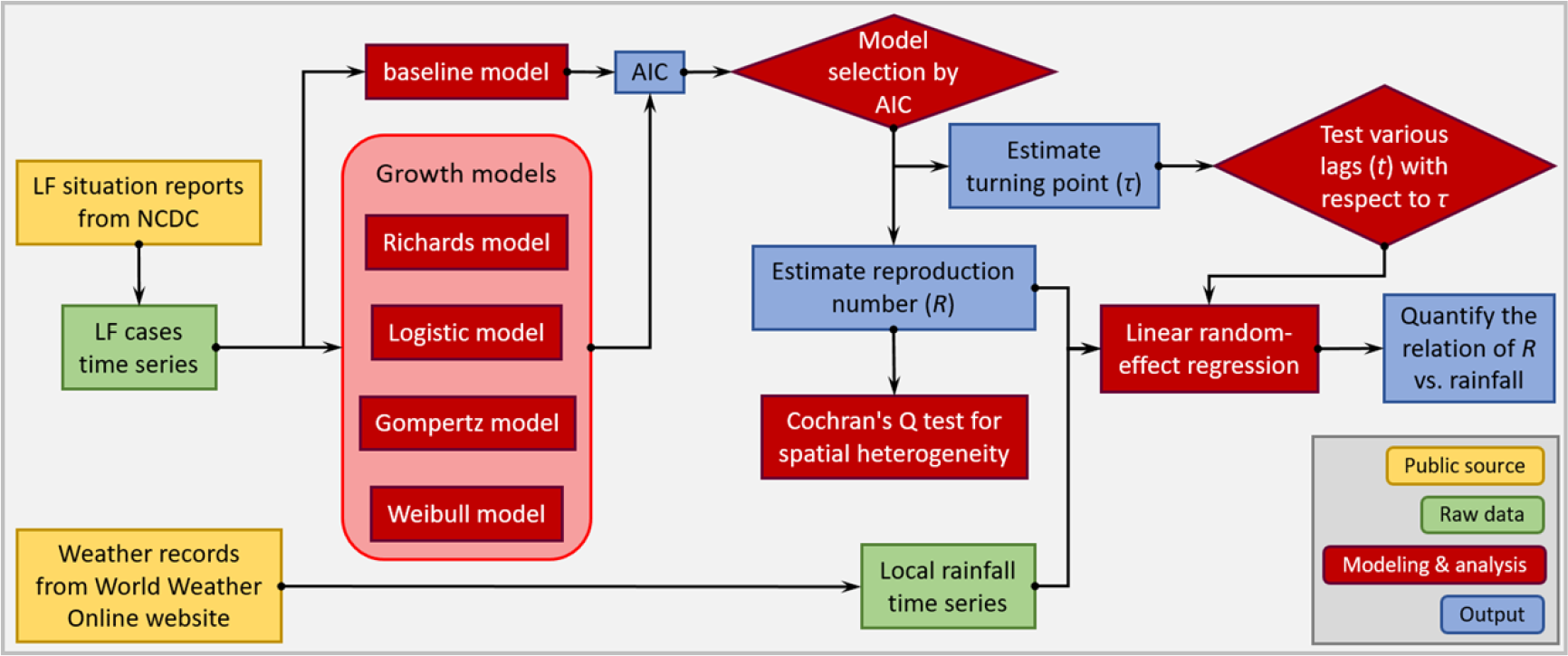
A flow diagram of the modeling analysis. This figure shows the analysis procedures in this study.

## Results and discussion

The rainfall time series of the five states and the weekly reported LF cases of the whole of Nigeria are shown in Figure 1(a)-(b). We observe that the major LF epidemics usually occur in Nigeria between November and May of the following year. The cumulative lagged effects can be observed by matching the peak timing of the rainfall and epidemic curves. In Figure 1(c), we shift the rainfall time series of the five states by +6.5 months to match the trends of the national LF epidemic curve in Nigeria. To further test the credibility of this match, we use a simple statistical regression model of “case ~ *α* exp(*β* * rainfall) + *θ*”, where *α*, *β* and *θ* are free parameters to be estimated, to check the outcome of the least-square fit. In Figure 1(d)-(e), we find that the fit has a *p*-value less than 0.0001, which indicates a significant (statistical) association between the LF cases and shifted rainfall curve.

We fit four different growth models to the LF confirmed cases and estimate the model-average reproduction number (*R*) after model selection. We show the growth model fitting results in Figure 3 and the model estimation and selection results in Table 1. The models fit the epidemic data well, and most of them have AICs lower than the baseline AR(2) model. Most of the regions exhibit an epidemic turning point (*τ*) ranging from epidemiological week (EW) 4 to 10 in each year. Out of four outbreaks in the states of Bauchi and Plateau, there are three estimated *τ*s after EW 10 (Table 1). The estimated reproduction number (*R*) of the outbreaks in different regions varies from 1.06 to 1.62. At the national level, the *R* value for the whole of Nigeria in 2016-17 (*R*=1.23 with 95% CI: [1.22, 1.24]) is significantly lower than in the epidemics of 2017-18 (*R*=1.33 with 95% CI: [1.29, 1.37]) and 2018-19 (*R*=1.29 with 95% CI: [1.27, 1.32]). The state of Edo has the highest estimated *R* (1.62 in 2017-18 and 1.45 in 2018-19), and this state also has the largest number of LF cases in the outbreaks of both 2017-18 (41.9% of all cases) and 2018-19 (36.0% of all cases). Hypothesized spatial heterogeneity in the *R* is tested by Cochran’s Q test. The testing results for the *R*s in the five states are significant (i.e., *p*-value < 0.05) for both the 2017-18 and 2018-19 LF epidemics. Thus, we report the existence of spatial heterogeneity in LF epidemics in Nigeria.

**Table 1.** The summary table of the model estimations. Population numbers are summarized in units of one million. The “improvement” is the goodness-of-fit estimate, i.e., the partial R-squared, from the baseline AR(2) model. The “weight” is the AIC-weight of the selected model, which is used for calculating the model-averaged estimates. The “NA” means a summary term that is not applicable to a certain model. The notation “-” means that the model cannot achieve a converging fitting outcome. The model-averaged estimates in each region are highlighted in gray.

| region | population | model | epidemic period | final size | reproduction number | turning point | R-squared | improvement | AIC | weight |
| --- | --- | --- | --- | --- | --- | --- | --- | --- | --- | --- |
| Nigeria | 186 | baseline: AR(2) | from EW44-2016 to EW22-2017 | NA | NA | NA | 0.9959 | 0.00% | 198.8 | NA |
| Nigeria | 186 | Richards | from EW44-2016 to EW22-2017 | - | - | - | - | - | - | - |
| Nigeria | 186 | Logistic | from EW44-2016 to EW22-2017 | 196; [191, 201] | 1.24; [1.22, 1.25] | EW10-2017; [EW10-2017, EW10-2017] | 0.9973 | 34.80% | 191.8 | 0.968 |
| Nigeria | 186 | Gompertz | from EW44-2016 to EW22-2017 | 221; [208, 233] | 1.14; [1.12, 1.15] | EW08-2017; [EW08-2017, EW09-2017] | 0.9969 | 23.90% | 198.6 | 0.032 |
| Nigeria | 186 | Weibull | from EW44-2016 to EW22-2017 | 350; [252, 448] | 1.05; [1.04, 1.07] | EW14-2017; [EW09-2017, EW18-2017] | 0.9937 | -53.80% | 220.5 | excluded |
| Nigeria | 186 | model average | from EW44-2016 to EW22-2017 | 197; [192, 202] | 1.23; [1.22, 1.24] | EW10-2017; [EW10-2017, EW10-2017] | NA | NA | NA | NA |
| Nigeria | 190.9 | baseline: AR(2) | from EW44-2017 to EW22-2018 | NA | NA | NA | 0.9977 | 0.00% | 157 | NA |
| Nigeria | 190.9 | Richards | from EW44-2017 to EW22-2018 | 464; [458, 470] | 1.33; [1.28, 1.37] | EW09-2018; [EW08-2018, EW09-2018] | 0.9984 | 32.00% | 151.2 | 0.933 |
| Nigeria | 190.9 | Logistic | from EW44-2017 to EW22-2018 | 468; [462, 474] | 1.41; [1.39, 1.43] | EW08-2018; [EW08-2018, EW08-2018] | 0.998 | 14.00% | 156.5 | 0.067 |
| Nigeria | 190.9 | Gompertz | from EW44-2017 to EW22-2018 | 474; [465, 483] | 1.31; [1.29, 1.34] | EW07-2018; [EW07-2018, EW07-2018] | 0.9972 | -20.60% | 169 | excluded |
| Nigeria | 190.9 | Weibull | from EW44-2017 to EW22-2018 | 491; [475, 507] | 1.08; [1.08, 1.08] | EW07-2018; [EW07-2018, EW07-2018] | 0.9955 | -91.10% | 183.3 | excluded |
| Nigeria | 190.9 | model average | from EW44-2017 to EW22-2018 | 464; [458, 470] | 1.33; [1.29, 1.37] | EW09-2018; [EW08-2018, EW09-2018] | NA | NA | NA | NA |
| Nigeria | 195.9 | baseline: AR(2) | from EW44-2018 to EW07-2019 | NA | NA | NA | 0.9959 | 0.00% | 66.8 | NA |
| Nigeria | 195.9 | Richards | from EW44-2018 to EW07-2019 | 447; [425, 469] | 1.29; [1.27, 1.32] | EW06-2019; [EW05-2019, EW06-2019] | 0.9982 | 56.10% | 57.5 | 1 |
| Nigeria | 195.9 | Logistic | from EW44-2018 to EW07-2019 | 596; [480, 713] | 1.39; [1.33, 1.45] | EW06-2019; [EW05-2019, EW07-2019] | 0.9924 | -87.60% | 78.7 | excluded |
| Nigeria | 195.9 | Gompertz | from EW44-2018 to EW07-2019 | 715; [431, 998] | 1.24; [1.15, 1.33] | EW05-2019; [EW03-2019, EW07-2019] | 0.991 | -121.60% | 83.4 | excluded |
| Nigeria | 195.9 | Weibull | from EW44-2018 to EW07-2019 | - | - | - | - | - | - | - |
| Nigeria | 195.9 | model average | from EW44-2018 to EW07-2019 | 447; [425, 469] | 1.29; [1.27, 1.32] | EW06-2019; [EW05-2019, EW06-2019] | NA | NA | NA | NA |
| Edo state | 4.2 | baseline: AR(2) | from EW50-2017 to EW18-2018 | NA | NA | NA | 0.9901 | 0.00% | 127.7 | NA |
| Edo state | 4.2 | Richards | from EW50-2017 to EW18-2018 | - | - | - | - | - | - | - |
| Edo state | 4.2 | Logistic | from EW50-2017 to EW18-2018 | 175; [173, 178] | 1.62; [1.58, 1.66] | EW09-2018; [EW08-2018, EW09-2018] | 0.9984 | 83.70% | 95.6 | 0.995 |
| Edo state | 4.2 | Gompertz | from EW50-2017 to EW18-2018 | 178; [174, 181] | 1.46; [1.42, 1.50] | EW08-2018; [EW08-2018, EW08-2018] | 0.9976 | 75.40% | 106.2 | 0.005 |
| Edo state | 4.2 | Weibull | from EW50-2017 to EW18-2018 | 182; [176, 189] | 1.13; [1.13, 1.13] | EW08-2018; [EW08-2018, EW08-2018] | 0.9963 | 62.20% | 115.3 | 0 |
| Edo state | 4.2 | model average | from EW50-2017 to EW18-2018 | 175; [173, 178] | 1.62; [1.58, 1.66] | EW09-2018; [EW08-2018, EW09-2018] | NA | NA | NA | NA |
| Ondo state | 4.7 | baseline: AR(2) | from EW48-2017 to EW18-2018 | NA | NA | NA | 0.9917 | 0.00% | 129.2 | NA |
| Ondo state | 4.7 | Richards | from EW48-2017 to EW18-2018 | - | - | - | - | - | - | - |
| Ondo state | 4.7 | Logistic | from EW48-2017 to EW18-2018 | 111; [108, 113] | 1.41; [1.37, 1.45] | EW07-2018; [EW06-2018, EW07-2018] | 0.9946 | 35.40% | 124.6 | 0.007 |
| Ondo state | 4.7 | Gompertz | from EW48-2017 to EW18-2018 | 114; [111, 117] | 1.28; [1.26, 1.31] | EW05-2018; [EW05-2018, EW06-2018] | 0.9968 | 61.50% | 114.7 | 0.99 |
| Ondo state | 4.7 | Weibull | from EW48-2017 to EW18-2018 | 128; [118, 138] | 1.14; [1.13, 1.15] | EW06-2018; [EW05-2018, EW06-2018] | 0.9947 | 36.40% | 126.3 | 0.003 |
| Ondo state | 4.7 | model average | from EW48-2017 to EW18-2018 | 114; [111, 117] | 1.28; [1.26, 1.31] | EW05-2018; [EW05-2018, EW06-2018] | NA | NA | NA | NA |
| Ebonyi state | 2.9 | baseline: AR(2) | from EW45-2017 to EW18-2018 | NA | NA | NA | 0.9937 | 0.00% | 117.8 | NA |
| Ebonyi state | 2.9 | Richards | from EW45-2017 to EW18-2018 | - | - | - | - | - | - | - |
| Ebonyi state | 2.9 | Logistic | from EW45-2017 to EW18-2018 | - | - | - | - | - | - | - |
| Ebonyi state | 2.9 | Gompertz | from EW45-2017 to EW18-2018 | - | - | - | - | - | - | - |
| Ebonyi state | 2.9 | Weibull | from EW45-2017 to EW18-2018 | 66; [64, 69] | 1.09; [1.09, 1.09] | EW07-2018; [EW07-2018, EW07-2018] | 0.997 | 52.60% | 104.6 | 1 |
| Ebonyi state | 2.9 | model average | from EW45-2017 to EW18-2018 | 66; [64, 69] | 1.09; [1.09, 1.09] | EW07-2018; [EW07-2018, EW07-2018] | NA | NA | NA | NA |
| Bauchi state | 6.5 | baseline: AR(2) | from EW49-2017 to EW16-2018 | NA | NA | NA | 0.9734 | 0.00% | 50.2 | NA |
| Bauchi state | 6.5 | Richards | from EW49-2017 to EW16-2018 | - | - | - | - | - | - | - |
| Bauchi state | 6.5 | Logistic | from EW49-2017 to EW16-2018 | - | - | - | - | - | - | - |
| Bauchi state | 6.5 | Gompertz | from EW49-2017 to EW16-2018 | - | - | - | - | - | - | - |
| Bauchi state | 6.5 | Weibull | from EW49-2017 to EW16-2018 | 17; [12, 21] | 1.09; [1.08, 1.09] | EW12-2018; [EW10-2018, EW13-2018] | 0.9889 | 58.40% | 37.5 | 1 |
| Bauchi state | 6.5 | model average | from EW49-2017 to EW16-2018 | 17; [12, 21] | 1.09; [1.08, 1.09] | EW12-2018; [EW10-2018, EW13-2018] | NA | NA | NA | NA |
| Plateau state | 4.2 | baseline: AR(2) | from EW48-2017 to EW16-2018 | NA | NA | NA | 0.9576 | 0.00% | 45.1 | NA |
| Plateau state | 4.2 | Richards | from EW48-2017 to EW16-2018 | - | - | - | - | - | - | - |
| Plateau state | 4.2 | Logistic | from EW48-2017 to EW16-2018 | - | - | - | - | - | - | - |
| Plateau state | 4.2 | Gompertz | from EW48-2017 to EW16-2018 | - | - | - | - | - | - | - |
| Plateau state | 4.2 | Weibull | from EW48-2017 to EW16-2018 | 16; [6, 25] | 1.07; [1.05, 1.09] | EW14-2018; [EW09-2018, EW19-2018] | 0.9872 | 69.70% | 24.7 | 1 |
| Plateau state | 4.2 | model average | from EW48-2017 to EW16-2018 | 16; [6, 25] | 1.07; [1.05, 1.09] | EW14-2018; [EW09-2018, EW19-2018] | NA | NA | NA | NA |
| Edo state | 4.3 | baseline: AR(2) | from EW46-2018 to EW08-2019 | NA | NA | NA | 0.9849 | 0.00% | 88.6 | NA |
| Edo state | 4.3 | Richards | from EW46-2018 to EW08-2019 | 140; [132, 149] | 1.43; [1.34, 1.51] | EW05-2019; [EW05-2019, EW06-2019] | 0.9972 | 81.20% | 71.1 | 0.856 |
| Edo state | 4.3 | Logistic | from EW46-2018 to EW08-2019 | 148; [139, 158] | 1.56; [1.50, 1.62] | EW05-2019; [EW05-2019, EW05-2019] | 0.9958 | 72.50% | 74.8 | 0.132 |
| Edo state | 4.3 | Gompertz | from EW46-2018 to EW08-2019 | 157; [140, 175] | 1.40; [1.32, 1.48] | EW04-2019; [EW04-2019, EW05-2019] | 0.995 | 66.90% | 79.6 | 0.012 |
| Edo state | 4.3 | Weibull | from EW46-2018 to EW08-2019 | - | - | - | - | - | - | - |
| Edo state | 4.3 | model average | from EW46-2018 to EW08-2019 | 142; [133, 150] | 1.45; [1.36, 1.53] | EW05-2019; [EW05-2019, EW06-2019] | NA | NA | NA | NA |
| Ondo state | 4.8 | baseline: AR(2) | from EW44-2018 to EW08-2019 | NA | NA | NA | 0.9913 | 0.00% | 80.3 | NA |
| Ondo state | 4.8 | Richards | from EW44-2018 to EW08-2019 | 133; [128, 137] | 1.27; [1.25, 1.29] | EW06-2019; [EW05-2019, EW07-2019] | 0.997 | 65.10% | 68.1 | 1 |
| Ondo state | 4.8 | Logistic | from EW44-2018 to EW08-2019 | 164; [134, 193] | 1.39; [1.31, 1.47] | EW05-2019; [EW04-2019, EW06-2019] | 0.9845 | -77.40% | 93.7 | excluded |
| Ondo state | 4.8 | Gompertz | from EW44-2018 to EW08-2019 | 171; [123, 218] | 1.30; [1.17, 1.42] | EW04-2019; [EW03-2019, EW06-2019] | 0.9814 | -113.20% | 98.9 | excluded |
| Ondo state | 4.8 | Weibull | from EW44-2018 to EW08-2019 | - | - | - | - | - | - | - |
| Ondo state | 4.8 | model average | from EW44-2018 to EW08-2019 | 133; [128, 137] | 1.27; [1.25, 1.29] | EW06-2019; [EW05-2019, EW07-2019] | NA | NA | NA | NA |
| Ebonyi state | 3 | baseline: AR(2) | from EW48-2018 to EW08-2019 | NA | NA | NA | 0.9627 | 0.00% | 59.1 | NA |
| Ebonyi state | 3 | Richards | from EW48-2018 to EW08-2019 | - | - | - | - | - | - | - |
| Ebonyi state | 3 | Logistic | from EW48-2018 to EW08-2019 | - | - | - | - | - | - | - |
| Ebonyi state | 3 | Gompertz | from EW48-2018 to EW08-2019 | - | - | - | - | - | - | - |
| Ebonyi state | 3 | Weibull | from EW48-2018 to EW08-2019 | 33; [24, 41] | 1.13; [1.12, 1.14] | EW05-2019; [EW04-2019, EW06-2019] | 0.99 | 73.20% | 48.5 | 1 |
| Ebonyi state | 3 | model average | from EW48-2018 to EW08-2019 | 33; [24, 41] | 1.13; [1.12, 1.14] | EW05-2019; [EW04-2019, EW06-2019] | NA | NA | NA | NA |
| Bauchi state | 6.7 | baseline: AR(2) | from EW48-2018 to EW08-2019 | NA | NA | NA | 0.9315 | 0.00% | 57.8 | NA |
| Bauchi state | 6.7 | Richards | from EW48-2018 to EW08-2019 | - | - | - | - | - | - | - |
| Bauchi state | 6.7 | Logistic | from EW48-2018 to EW08-2019 | - | - | - | - | - | - | - |
| Bauchi state | 6.7 | Gompertz | from EW48-2018 to EW08-2019 | - | - | - | - | - | - | - |
| Bauchi state | 6.7 | Weibull | from EW48-2018 to EW08-2019 | 141; [0, 474] | 1.06; [1.02, 1.10] | EW16-2019; [EW02-2019, EW02-2020] | 0.9819 | 73.60% | 47.3 | 1 |
| Bauchi state | 6.7 | model average | from EW48-2018 to EW08-2019 | 141; [0, 474] | 1.06; [1.02, 1.10] | EW16-2019; [EW02-2019, EW02-2020] | NA | NA | NA | NA |
| Plateau state | 4.3 | baseline: AR(2) | from EW48-2018 to EW08-2019 | NA | NA | NA | 0.9467 | 0.00% | 56.4 | NA |
| Plateau state | 4.3 | Richards | from EW48-2018 to EW08-2019 | 33; [30, 37] | 1.35; [1.24, 1.44] | EW06-2019; [EW05-2019, EW07-2019] | 0.9873 | 76.10% | 44.4 | 0.64 |
| Plateau state | 4.3 | Logistic | from EW48-2018 to EW08-2019 | 37; [31, 43] | 1.50; [1.37, 1.62] | EW05-2019; [EW04-2019, EW06-2019] | 0.9804 | 63.20% | 48.1 | 0.104 |
| Plateau state | 4.3 | Gompertz | from EW48-2018 to EW08-2019 | 35; [30, 40] | 1.54; [1.34, 1.72] | EW05-2019; [EW04-2019, EW05-2019] | 0.9841 | 70.20% | 47.3 | 0.15 |
| Plateau state | 4.3 | Weibull | from EW48-2018 to EW08-2019 | 37; [29, 45] | 1.14; [1.13, 1.15] | EW05-2019; [EW04-2019, EW05-2019] | 0.9832 | 68.50% | 48 | 0.106 |
| Plateau state | 4.3 | model average | from EW48-2018 to EW08-2019 | 34; [30, 39] | 1.37; [1.26, 1.47] | EW05-2019; [EW04-2019, EW06-2019] | NA | NA | NA | NA |

**Figure 3.**
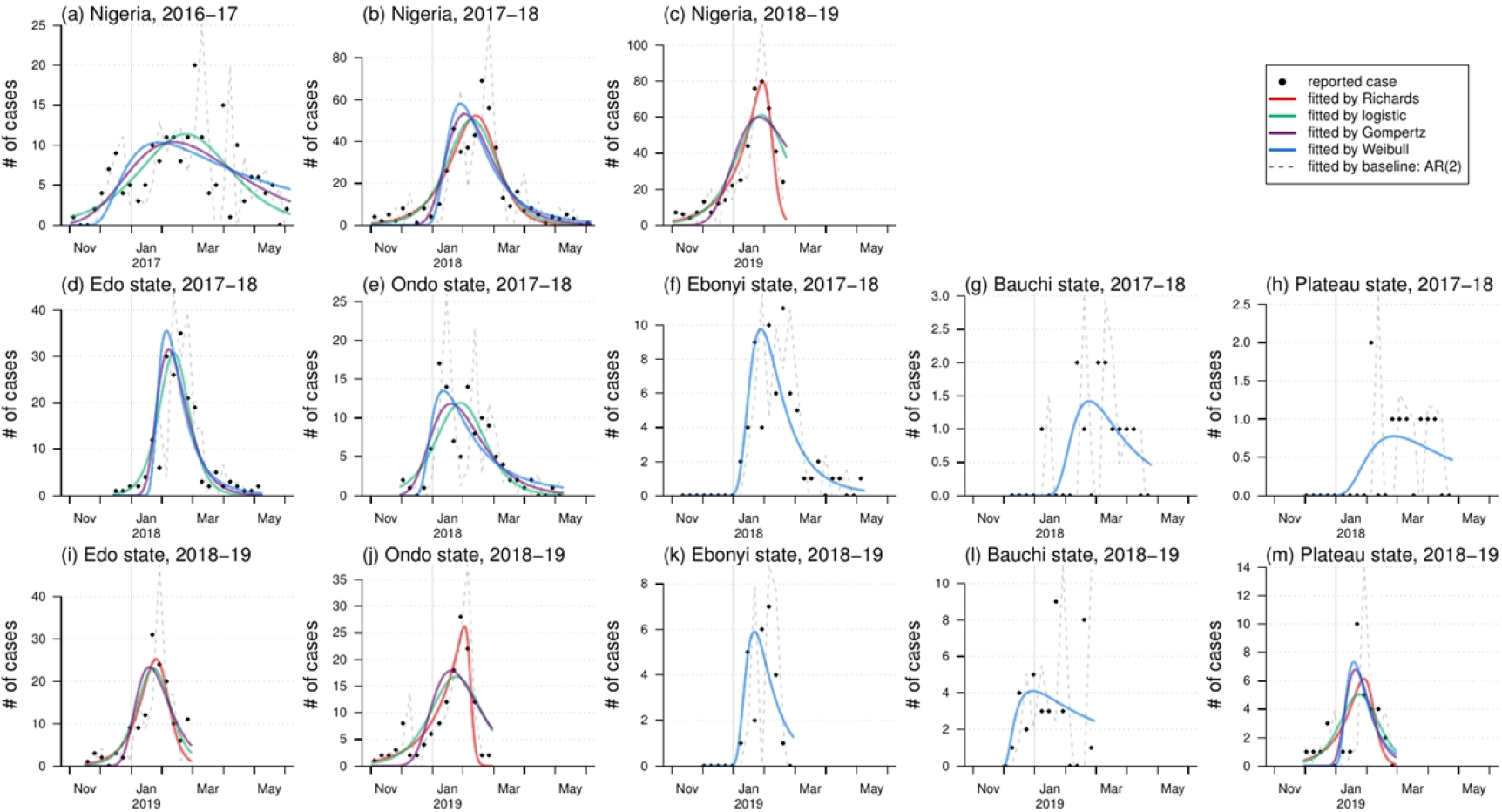
The fitting results of estimates of Lassa epidemics in Nigeria by nonlinear growth models. In each panel, the dots are the observed (reported) cases, the dashed gray line is the fit by the baseline AR(2) model, and the colored lines are the fits from the nonlinear growth models.

We choose to use the average reproduction number (*R*) rather than the instantaneous reproduction number, commonly denoted by *R*_*t*_ [22, 23], as the measurement of the LASV transmissibility. The *R*_*t*_ is a series of time-dependent reproduction numbers, namely, time-dependent effective reproduction numbers, which can be estimated by a renewable equation [20, 22, 23, 33, 34]. The factors that affect the changing dynamics of *R_t,_* include (i) the depletion of the susceptible population [28], (ii) the change (usually it is the improvement) in the unmeasurable disease control efforts (e.g., contract tracing) and local awareness of the outbreak [29], and (iii) the natural features of the pathogen (e.g., its original infectivity and other interepidemic factors) [28, 29, 31]. The estimated *R* in this work is a point estimate to summarize LASV transmissibility over a whole epidemic. Hence, the temporal changes of the susceptible population in (i), above, and local disease awareness and control related to (ii) do not affect *R*. With respect to point (iii) and other heterogeneities of outbreaks in different regions, we account for this issue by including the “region” dummy variables in the LRER model in Eqn (3). These dummy variables serve as random effects to offset regional heterogeneities in LF epidemics. Therefore, we can then quantify a consistent effect (the *β* in Eqn (3)) of the lagged rainfall on the LASV *R* estimate.

The association between local rainfall and LASV transmissibility (*R*) are modeled and quantified by the LRER model. In Figure 4, we find a positive relation between rainfall and *R*. The estimated change rate in *R* for a one unit (mm) increase in rainfall is summarized with different cumulative lag terms (the *t* in Eqn (3)).

- For a cumulative lag of 4 months, the change rate is 0.42% (95% CI: [0.02%, 0.82%]; *p*-value: 0.039) with a regression R-squared of 0.36.
- For a cumulative lag of 5 months, the change rate is 0.45% (95% CI: [0.04%, 0.87%]; *p*-value: 0.034) with a regression R-squared of 0.37.
- For a cumulative lag of 6 months, the change rate is 0.62% (95% CI: [0.15%, 1.08%]; *p*-value: 0.012) with a regression R-squared of 0.44.
- For a cumulative lag of 7 months, the change rate is 0.62% (95% CI: [0.20%, 1.05%]; *p*-value: 0.006) with a regression R-squared of 0.47.
- For a cumulative lag of 8 months, the change rate is 0.57% (95% CI: [0.12%, 1.02%]; *p*-value: 0.016) with a regression R-squared of 0.42.
- For a cumulative lag of 9 months, the change rate is 0.54% (95% CI: [0.36%, 1.06%]; *p*-value: 0.044) with a regression R-squared of 0.36.

**Figure 4.**
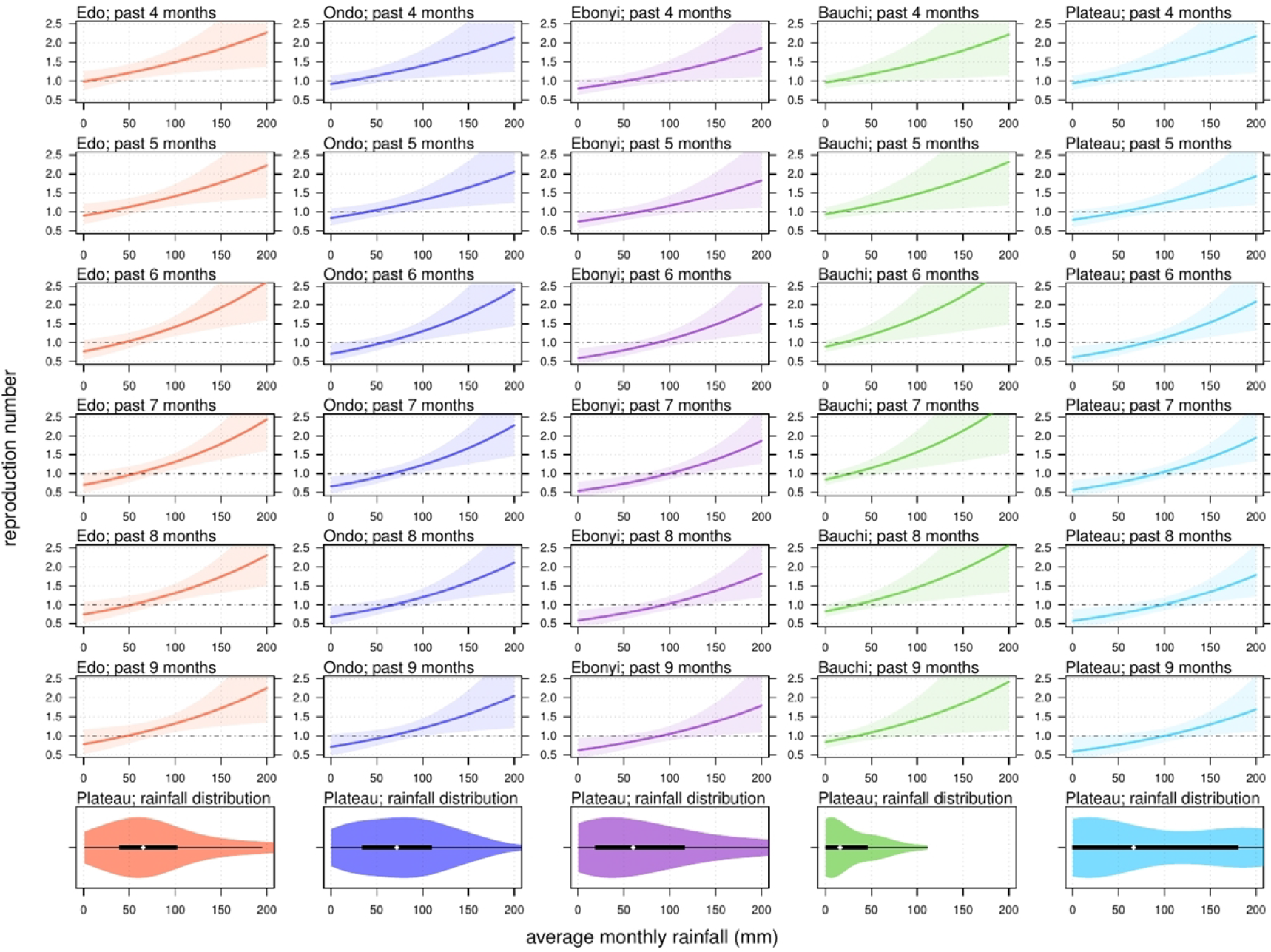
The relationship between local rainfall and Lassa fever (LF) transmissibility, i.e., the reproduction number (*R*), in five different states with different time lags (*t*). The reproduction number of 1.0 is highlighted by the back dashed line. The panels at the bottom are the violin plots and show the distribution of local rainfall in each state. The black rectangles represent the 25% and 75% quantiles, and the white dot is the median.

We report the most significant (i.e., with the lowest *p*-value) regression estimates that appear with a cumulative lag of 7 months. The habitats of the LASV reservoir, i.e., rodents, include irrigated and flooded agricultural lands that are commonly found in and around African villages [6]. The range of lag from 4-9 months has previously been explained by the time interval between the peak of the rainfall and the rodent population [7]. The association between rodent population dynamics and rainfall levels has been demonstrated in a number of previous studies [6–10]. This relation could also be verified by examining the rodent population data in the Nigerian regions included in this work. The present finding of the impact of lagged rainfall on LF outbreaks suggests that knowledge of such weather-driven epidemics could be gained by referring to past rainfall levels. For instance, if a relatively high amount of rainfall occurs, local measures, such as rodent population control, could be effective to reduce the LF risk. The findings in this work are of public health interest and are helpful for policy-makers trying to achieve LF prevention and control.

On the one hand, our findings suggest the existence of an association between rainfall and LASV transmissibility, which could be affected by the population dynamics of rodents [13]. On the other hand, the positive relation between rainfall and *R* indicates that rainfall, particularly in states with a high LF risk, can be used as a warning signal for LF epidemics. The modeling framework in this study can be easily extended to other infectious diseases.

## Conclusions

The LF epidemic reproduction numbers (*R*) of the whole of Nigeria in 2017-18 (*R*=1.33 with 95% CI: [1.29, 1.37]) and 2018-19 (*R*=1.29 with 95% CI: [1.27, 1.32]) are significantly higher than in 2016-17 (*R*=1.23 with 95% CI: [1.22, 1.24]). There is significant spatial heterogeneity in the LF epidemics of different Nigerian regions. We report clear evidence of rainfall impacts on LF outbreaks in Nigeria and quantify the impact. A one unit (mm) increase in average rainfall over the past 7 months could cause a 0.62% (95% CI: [0.20%, 1.05%]) rise in the *R*. Local rainfall can be used as a warning signal for LF epidemics.

## List of abbreviations

LASV: Lassa virus
LF: Lassa fever
CI: confidence interval
EW: epidemiological week
MGF: moment generating function
SD: standard deviation
AIC: Akaike information criterion
NLS: nonlinear least squares
AR: autoregression
NCDC: Nigeria Centre for Disease Control
LRER: linear random-effect regression

## Declarations

### Ethics approval and consent to participate

Since no personal data were collected, ethical approval and individual consent were not applicable.

### Availability of data and material

All data used for analysis are freely available on online public domains [12, 15].

### Consent for publication

Not applicable.

## Acknowledgments

Not applicable.

## Disclaimer

The funding agencies had no role in the design and conduct of the study; collection, management, analysis, and interpretation of the data; preparation, review, or approval of the manuscript; or decision to submit the manuscript for publication.

## Authors’ Contributions

SZ conceived, carried out the study, and drafted the first manuscript. SZ and DH discussed the results. All authors revised the manuscript and gave final approval for publication.

